# Brain Iron Mediates the Relationship Between Cognition and Neighborhood Socioeconomic Status in Youth

**DOI:** 10.1101/261693

**Authors:** Lauren Beard, David R. Roalf, Mark A. Elliott, Karthik Prabhakaran, Tyler M. Moore, Raquel E. Gur, Theodore D. Satterthwaite

**Author notes:** **Please address correspondence to:** Theodore D. Satterthwaite, M.D., 10^th^ Floor, Gates Building, Hospital of the University of Pennsylvania, 34^th^ and Spruce Street, Philadelphia, PA 19104.

## Abstract

Non-heme brain iron is a critical metabolic cofactor essential for healthy brain development.^1^ Iron deficiency is the most common nutritional disorder in the world, with greater prevalence of non-heme iron deficiency among individuals of lower socioeconomic status (SES). However, it remains unknown how brain iron accumulation during development may impact cognition. Brain iron can be measured *in vivo* using R2* weighted magnetic resonance imaging (MRI); prior work has established that higher R2* is associated with higher iron content.^2,3^ We hypothesized that more iron in the basal ganglia (BG) regions of the caudate, putamen, and pallidum would be associated with improved cognitive performance and potentially mediate the known relationship between neighborhood-level socioeconomic status (SES) and cognition.^4^

## METHODS

We quantified BG iron in a large sample of youth ages 8-23 imaged as part of the Philadelphia Neurodevelopmental Cohort (PNC), a community-based study of brain development.^5^ In total, we considered 1,147 youth (mean age 14.90, SD = 3.61; 524 males and 623 females), after excluding those with missing data, poor quality imaging, or medical problems that could impact brain function. All participants completed the Penn Computerized Neurocognitive Battery (CNB); overall accuracy was summarized from the CNB using factor analysis.^6^ Socioeconomic status was quantified on the neighborhood level as in previous work from the PNC.^4^

All participants completed both structural imaging and a multi-echo sequence that allowed quantification of R2*. BG structures were delineated for each subject using an advanced multi-atlas labeling procedure. Mean R2* signal within each bilateral BG region was modeled using generalized additive models (GAM) with penalized splines to capture both linear and nonlinear age effects. We investigated main effects of age, sex, race, and cognition at each region while correcting for multiple comparisons using the False Discovery Rate (*q* <0.05); interactions were evaluated but found to be non-significant. Finally, we tested whether BG iron mediated the relationship between SES and cognition using a mediation analysis with bootstrapped confidence intervals. The potentially confounding influence of data quality was evaluated by adding the quality rating of the structural image as a model covariate. Specificity of neighborhood SES effects was examined by including maternal education as an additional model covariate. See https://goo.gl/dkVbaC for all analysis code.

## RESULTS

As expected from prior reports, R2* increased in the BG throughout youth (**Figure 1**; all regions *p*_*fdr*_ <0.001); no sex differences or age by sex interactions were present. Across all ages, lower R2* levels in the caudate (*p*_*fdr*_<0.02) was associated with diminished cognitive performance. Lower neighborhood SES was associated with both diminished cognition (*p*<0.001), as well as lower R2* in the caudate (*p*_*fdr*_=0.02). Even when controlling for maternal education, SES remained significantly associated with caudate R2* (*p*=0.03). Critically, the relationship between cognition and SES was significantly mediated by caudate R2* (**Figure 2**; *p*=0.007). Including data quality as a covariate did not impact results.

**Figure 1.**
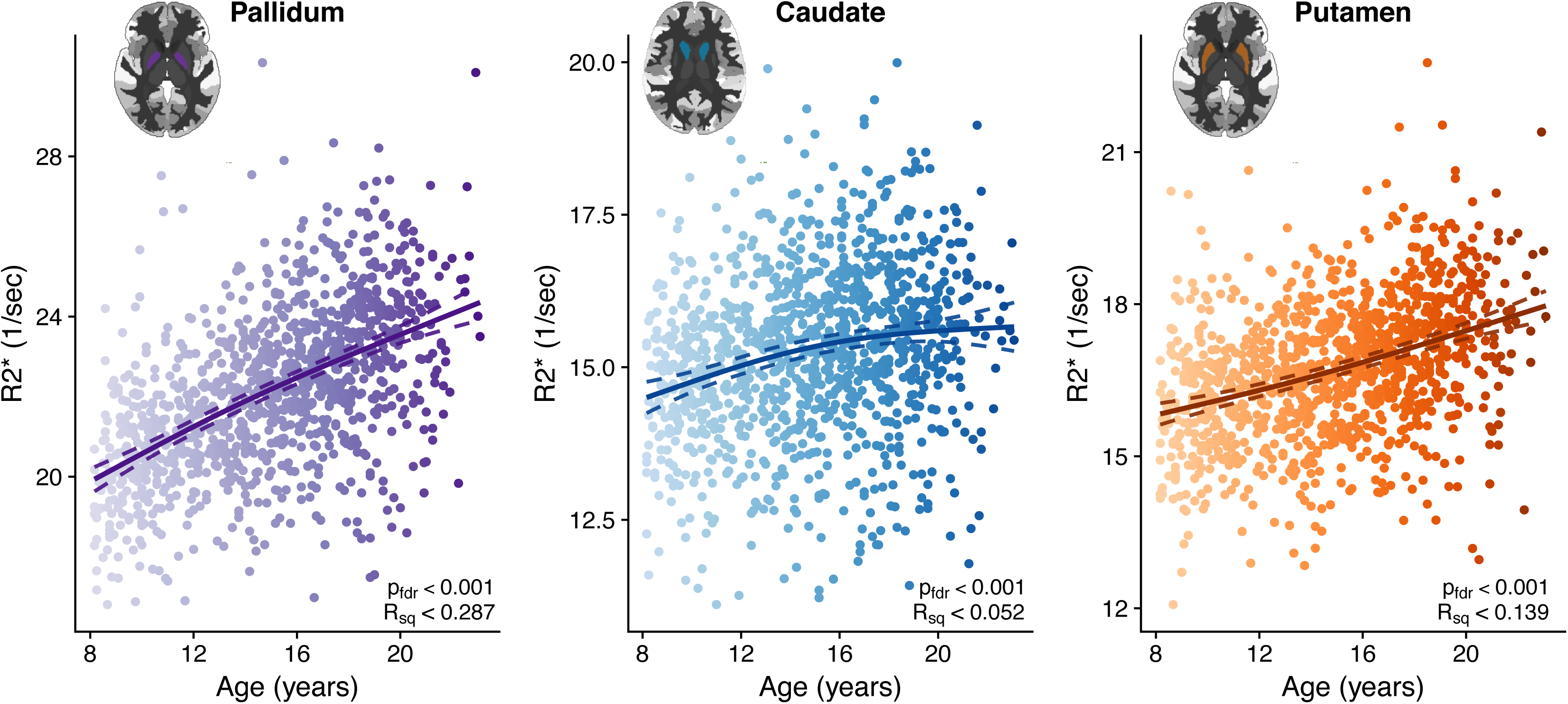
R2* indexes brain iron and accumulates during development. R2* accumulates in the brain throughout development in the pallidum (A), caudate (B), and putamen (C). No sex differences or age by sex interactions were present.

**Figure 2.**
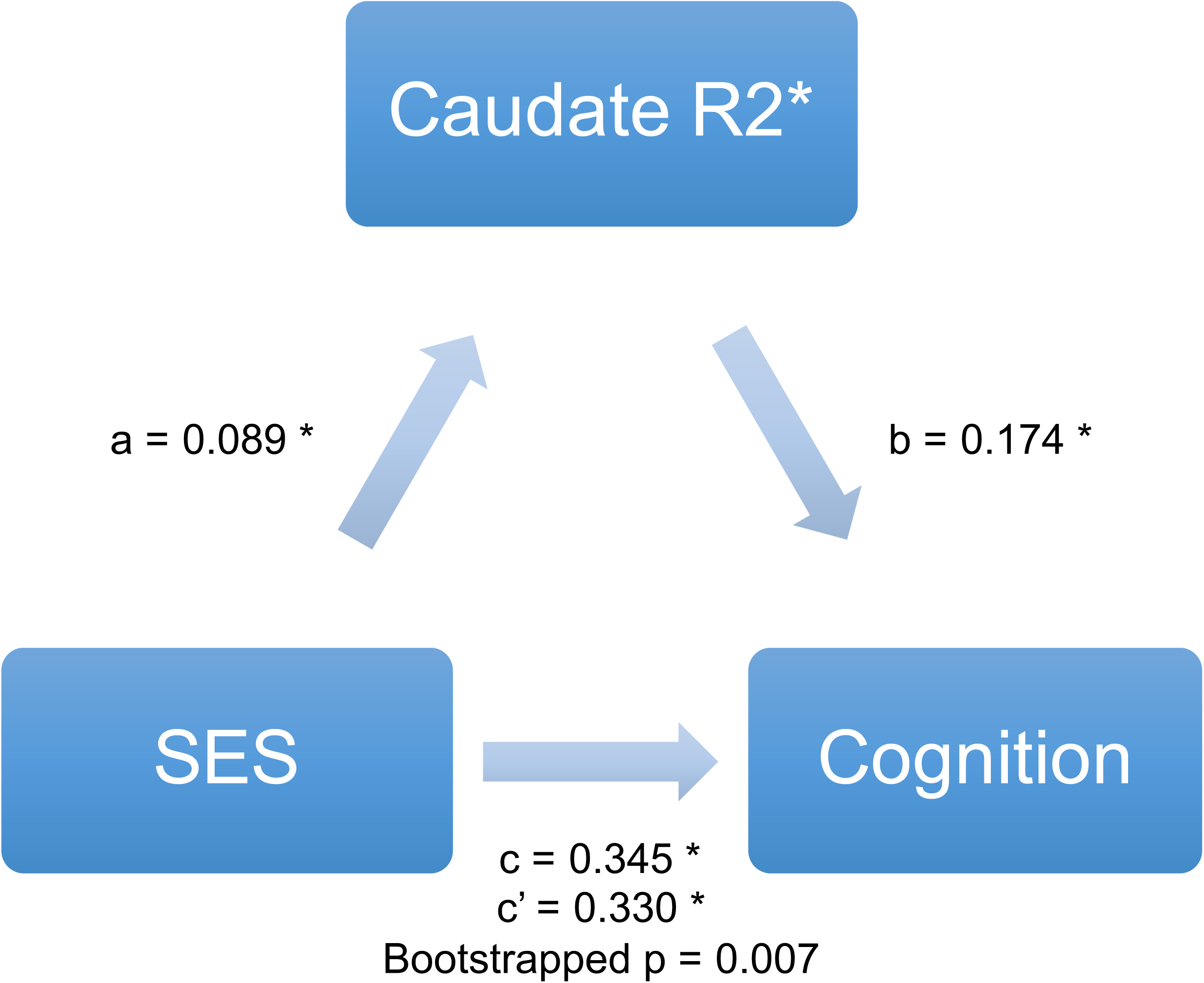
The relationship between SES and cognition is mediated by R2*. Mediation results are shown as standardized regression coefficients. Significance of indirect effect (ab = 0.018) was assessed using bootstrapped confidence intervals [0.005 − 0.028]. The asterisk (*) indicates p < 0.01.

## DISCUSSION

Leveraging a large sample of youth imaged as a part of the PNC, we replicate prior results documenting that brain iron indexed by R2* accumulates with age in the basal ganglia in youth.^1,3^ Additionally, we provide novel evidence that lower levels of caudate iron is associated with diminished cognitive performance during adolescence. Importantly, lower neighborhood-level SES is also associated with caudate brain iron, and brain iron significantly mediated the relationship between cognition and SES. While speculative, it is possible that these effects are due to neighborhood-level differences in nutrition and the availability of iron-rich foods. Taken together, these results suggest that brain iron is important for facilitating cognitive development in youth, and may provide a mechanism by which public health interventions focused on improving nutrition in low SES neighborhoods might aid cognitive development.

## AUTHOR CONTRIBUTIONS

Beard and Satterthwaite had full access to all of the data and take responsibility for the integrity of the data and accuracy of the analysis.*Study concept and design:* Beard, Roalf, Elliott, Prabhakaran, Satterthwaite, *Acquisition, analysis, or interpretation of data:* all authors *Drafting of the manuscript*: Beard, Roalf, Satterthwaite *Critical revision of the manuscript for important intellectual content*: all authors *Statistical analyses*: Beard, Elliott, Moore, Satterthwaite *Obtained funding*: Roalf, Gur RC, Gur RE, Satterthwaite *Administrative, technical, or material support*: Elliott, Prabhakaran

## ADDITIONAL CONTRIBUTIONS

Many thanks to Adon Rosen for image processing, to Monica Calkins for phenotyping expertise, to Russell T. Shinohara for statistical advice, and to Ruben Gur with cognitive assessment. Thanks to Chad Jackson and Kosha Ruparel for data management and systems support.

